# The African swine fever virus protease pS273R inhibits DNA sensing cGAS-STING pathway by targeting IKKε

**DOI:** 10.1101/2021.06.02.446855

**Authors:** Jia Luo, Jinghua Ni, Sen Jiang, Nengwen Xia, Yiwen Guo, Qi Shao, Jiajia Zhang, Qi Cao, Zhenqi Xu, Wanglong Zheng, Nanhua Chen, Quan Zhang, Hongjun Chen, Xiaoyu Guo, Hongfei Zhu, François Meurens, Jianzhong Zhu

## Abstract

African swine fever virus (ASFV), a large and complex cytoplasmic double-stranded DNA virus, has developed multiple strategies to evade the antiviral innate immune responses. Cytosolic DNA arising from invading ASFV is mainly detected by the cyclic GMP-AMP synthase (cGAS) and then triggers a series of innate immune responses to prevent virus invasion. However, the immune escape mechanism of ASFV remains to be fully clarified. The pS273R of ASFV is a member of the SUMO-1-specific protease family and is crucial for valid virus replication. In this study, we identified pS273R as a suppressor of cGAS-STING pathway mediated type I interferon (IFN) production by ASFV genomic open reading frame screening. The pS273R was further confirmed as an inhibitor of IFN production as well as its downstream antiviral genes in cGAS-STING pathway. Mechanistically, pS273R greatly decreased the cGAS-STING signaling by targeting IKKε but not TBK1 and pS273R was found to disturb the interaction between IKKε and STING through its interaction with IKKε. Further, mutational analyses revealed that pS273R antagonized the cGAS-STING pathway by enzyme catalytic activity, which may affect the IKKε sumoylation state required for the interaction with STING. In summary, our results revealed for the first time that pS273R acts as an obvious negative regulator of cGAS-STING pathway by targeting IKKε via its enzymatic activity, which shows a new immune evasion mechanism of ASFV.

**Importance:** African swine fever (ASF) is a devastating disease for domestic pigs and wild boar and the pathogen ASFV is a cytoplasmic double-stranded DNA virus. The innate immune cGAS-STING-IFN signaling pathway exerts a critical role in sensing ASFV infection. However, the functions of half ASFV encoded 150 plus proteins are still unknown and the evasion against the cGAS-STING pathway is not resolved. In our study, via ASFV genomic open reading frame (ORF) screening, we found that 29 ASFV proteins could inhibit cGAS-STING signaling pathway, with pS273R showing the most obvious inhibitory effect. Surprisingly, pS273R was found to antagonize the cGAS-STING signaling by targeting IKKε. Moreover, the pS273R enzyme activity is required for its ability to inhibit the cGAS-STING pathway. Our findings deepen the understanding of the immune evasion mechanism of ASFV, which will provide a support for the development of safe and effective ASFV vaccines.

## Introduction

African swine fever (ASF), a devastating disease for domestic pigs and wild boar, usually causes an acute hemorrhagic fatal disease with a mortality rate of up to 100% while being asymptomatic in the natural hosts (1, 2). Since African swine fever was initially described in Kenya in the 1920s, it was spread rapidly across many countries in sub-Saharan African, Caribbean, Eastern Europe (3). In 2017, a large number of outbreaks of ASF occurred in Siberia near to the Russia-China border (4). The first outbreak of ASF in China was reported on August 3, 2018 (5) where it caused unprecedented disaster for Chinese swine industry and food security due to currently unavailable vaccines in the world. Accordingly, animal slaughter and regional quarantine are the only method for the disease control.

African swine fever virus (ASFV) is a large, complex, cytoplasmic double-stranded DNA virus. It is the sole member of the *Asfarviridae* family, but also the only DNA virus transmitted by arthropod ticks, soft ticks (*Ornithodoros moubata*) (6). ASFV is an enveloped virus with icosahedral structure inside which is composed of four concentric layers from the central nucleoid, the core shell, the inner envelope to the outside the icosahedral capsid (7). The average diameter of the ASFV is approximately 200 nm and genomes vary in length between 170 and 193 kb which depends on the isolates. It contains between 151 and 167 open reading frames (ORFs), which encode structural proteins, viral DNA replication proteins and host defense escape protein *etc* (8).

The open reading frame (ORF) S273R encodes a 31-kD protein, and the protein pS273R belongs to the SUMO-1-specific protease family with the ability to catalyze the maturation of pp220 and pp62 polyprotein precursors into core shell matrix proteins (9). The two significant polyproteins, pp220 and pp62, are cleaved by the pS273R protease to produce six primary structural components of the virus particle, among which p37, p34, p40, p150 derived from polyproteins pp220 and p15, p35 derived from polyproteins pp62 (10). Therefore, the pS273R protease is of great significance for maturation and infectivity of the ASFV particle.

The innate immune system is the body’s first line of defense against pathogen infection. It utilizes a variety of pattern recognition receptors (PRRs) in cells to recognize and respond to pathogen associated molecular patterns (PAMPs) (11). Among PRRs, cyclic GMP-AMP synthase (cGAS) is a cytosolic double-stranded DNA sensor, which senses the presence of cytoplasmic DNA and catalyzes the synthesis of the second messenger cyclic GMP-AMP (2’3’-cGAMP) (12). Then 2’3’-cGAMP binds and activates the stimulator of interferon genes protein (STING), which upon stimulation, transfers from the endoplasmic reticulum (ER) to the trans-Golgi apparatus network (TGN), during which the kinase TBK1 is recruited and phosphorylated (13). TBK1 subsequently phosphorylates the transcription factor IRF3, and then IRF3 translocates into nucleus to activate the expression of antiviral type I interferons (IFNs) (14). The DNA sensing cGAS-STING pathway is the relevant innate immunity for ASFV, however, the immune evasion of this pathway by ASFV has not been resolved. Here, we identified the viral protein responsible for the immune evasion and characterized its mechanism of action.

## Results

### The ASFV pS273R protein significantly inhibits cGAS-STING signaling activity

The cGAS serves as an essential cytosolic DNA sensor that upon activation quickly triggers STING-dependent signaling and results in transcriptions of type I interferon (IFN) genes and proinflammatory cytokine genes (15). ASFV is a cytoplasmic dsDNA virus and mainly activates DNA-sensing cGAS-STING signaling pathway (16). Recently, two studies reported that ASFV pDP96R and pA528R (MGF505-7R) suppress cGAS-STING pathway mediated production of IFNs and proinflammatory cytokines by targeting TBK1/IKKβ and STING, respectively (17, 18). However, the question whether the cGAS-STING pathway is suppressed by other ASFV viral proteins still persist. In our study, we utilized the porcine cGAS-STING activated ISRE promoter assay and screened the ASFV genomic ORFs for modulators of cGAS-STING pathway (Supplementary Fig 1). The results showed that a number of ASFV genes exhibited an obvious inhibiting effect and were likely to play a role of immune evasion in the cGAS-STING pathway. These genes include MGF110-1L, MGF110-2L, MGF110-3L, MGF110-4L, MGF110-5-6L, MGF360-4L, X69R, MGF300-1L, MGF300-4L, MGF505-1R, MGF360-12L, MGF505-6R, MGF505-7R, A104R, A118R, A151R, F334L, F778R, EP152R, EP153R, EP402R, CP204L, CP80R, D345L, S183L, S273R, E120R, I215L and DP96R. Among these inhibitory genes, S273R encoding pS273R was the most effective and showed reproducible suppression of cGAS-STING signaling which was not reported previously. Therefore, we pursued to investigate the inhibitory role of S273R in the cGAS-STING pathway.

We first confirmed the inhibitory role of pS273R using different promotor assays in transfected 293T cells, and found that co-transfection of pS273R in 293T cells could significantly inhibit ISRE, IFNβ and NF-κB promoter activity induced by porcine cGAS-STING signaling (Fig 1A). Co-transfection of porcine cGAS and STING activated downstream IFNβ, ISG56, ISG60 and IL-8 gene transcriptions and the inductions of these gene transcriptions were significantly suppressed by the co-transfected pS273R (Fig 1B). Co-expression of porcine cGAS and STING led to the phosphorylation of TBK1 and IRF3, and downstream ISG56 production. The presence of pS273R retained the phosphorylation of TBK1, but restrained the phosphorylation of IRF3 and the production of ISG56 (Fig 1C).

**Figure 1.**
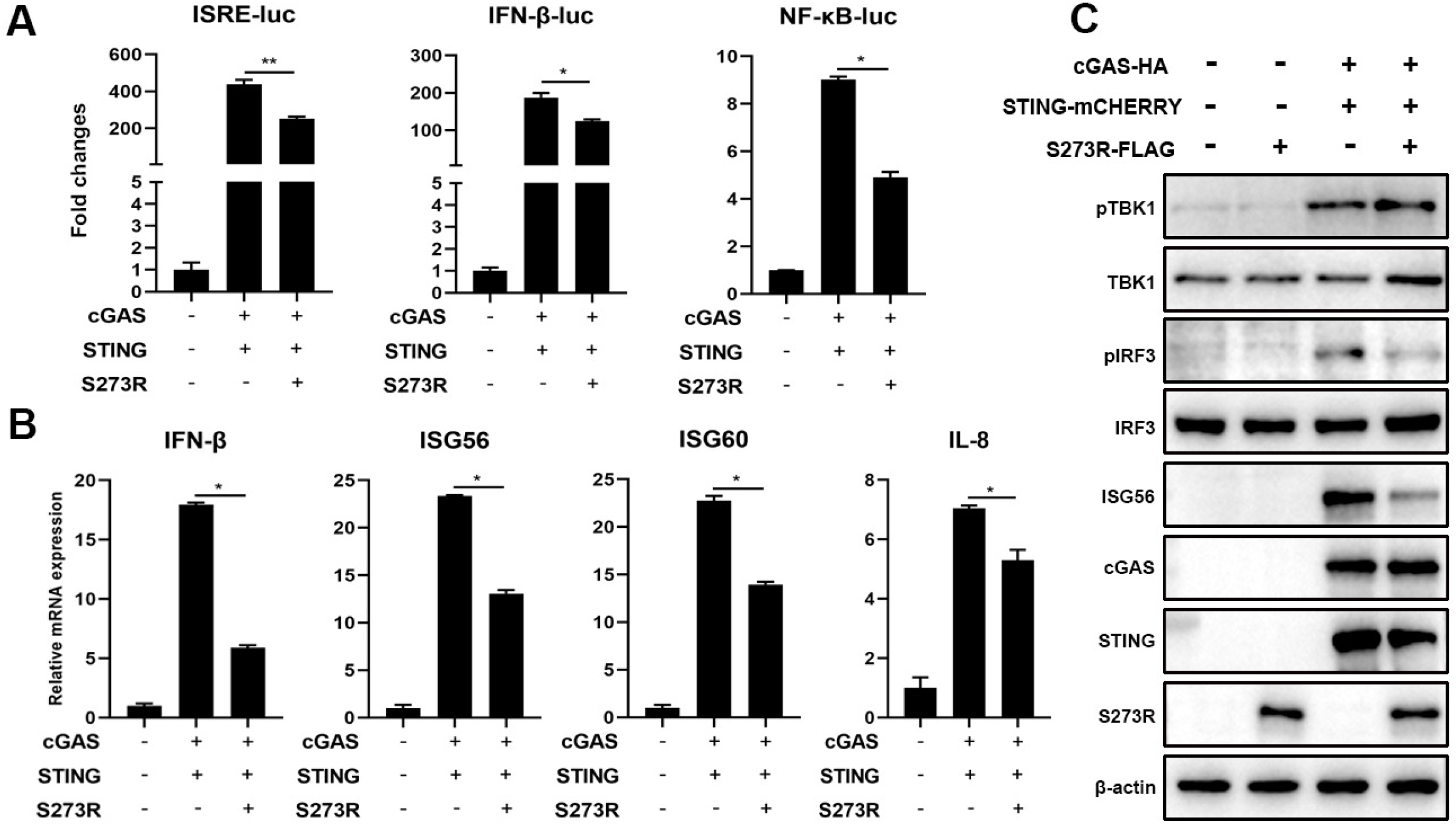
The effect of ASFV pS273R on the signaling activity of exogenous porcine cGAS-STING pathway. (A) HEK293T cells in 96-well plates (2×10^4^ cells/well) were co-transfected with 20 ng cGAS-HA and 10 ng STING-mCHERRY, plus 10 ng ISRE-luc or IFNβ-luc or NF-κβ-luc and 0.2 ng pRL-TK plasmid, along with 10 ng 3×FLAG-pCMV-S273R or control vector 3×FLAG-pCMV, which were normalized to 50 ng/well by control vector. At 24h post-transfection, luciferase activities were detected using Double-Luciferase Reporter Assay. (B-C) HEK293T cells in 24-well plates (3×10^5^ cells/well) were co-transfected with 400 ng cGAS-HA and 400 ng STING-mCHERRY plasmids together with 400 ng 3 × FLAG-pCMV-S273R or control vector for 24h, then the cells were harvested and analyzed by RT-qPCR for downstream gene expressions (B) and Western blotting using the indicated antibodies (C).

In order to explore the effect of pS273R on endogenous porcine cGAS-STING signaling pathway, we used the PAMs stimulated by three different agonists, polydA:dT and 45bp dsDNA to activate cGAS, and 2’3’-cGAMP to activate STING. In promoter assays, we found that pS273R significantly inhibited polydA:dT and 2’3’-cGAMP activated ISRE and NF-κB promoter activity (Fig 2A). In RT-qPCR assay, the downstream gene transcriptions of porcine IFNβ, ISG56, IL-8 mRNA levels induced by polydA:dT (Fig 2B), 45bp dsDNA (Fig 2C) and 2’3’-cGAMP (Fig 2D) were all significantly inhibited by pS273R. Consistently, we also found that phosphorylation of IRF3 and expression of ISG56 were markedly decreased in PAMs in the presence of pS273R after polydA:dT stimulation (Fig 2E) and 2’3’-cGAMP stimulation (Fig 2F). However, in both cases, the phosphorylation of TBK1 was not affected by pS273R (Fig 2E and 2F). These results were perfectly in line with those from exogenous porcine cGAS-STING signaling pathway in transfected 293T cells (Fig 1). Taken together, our results clearly suggested that pS273R inhibits porcine DNA sensing cGAS-STING signaling activity.

**Figure 2.**
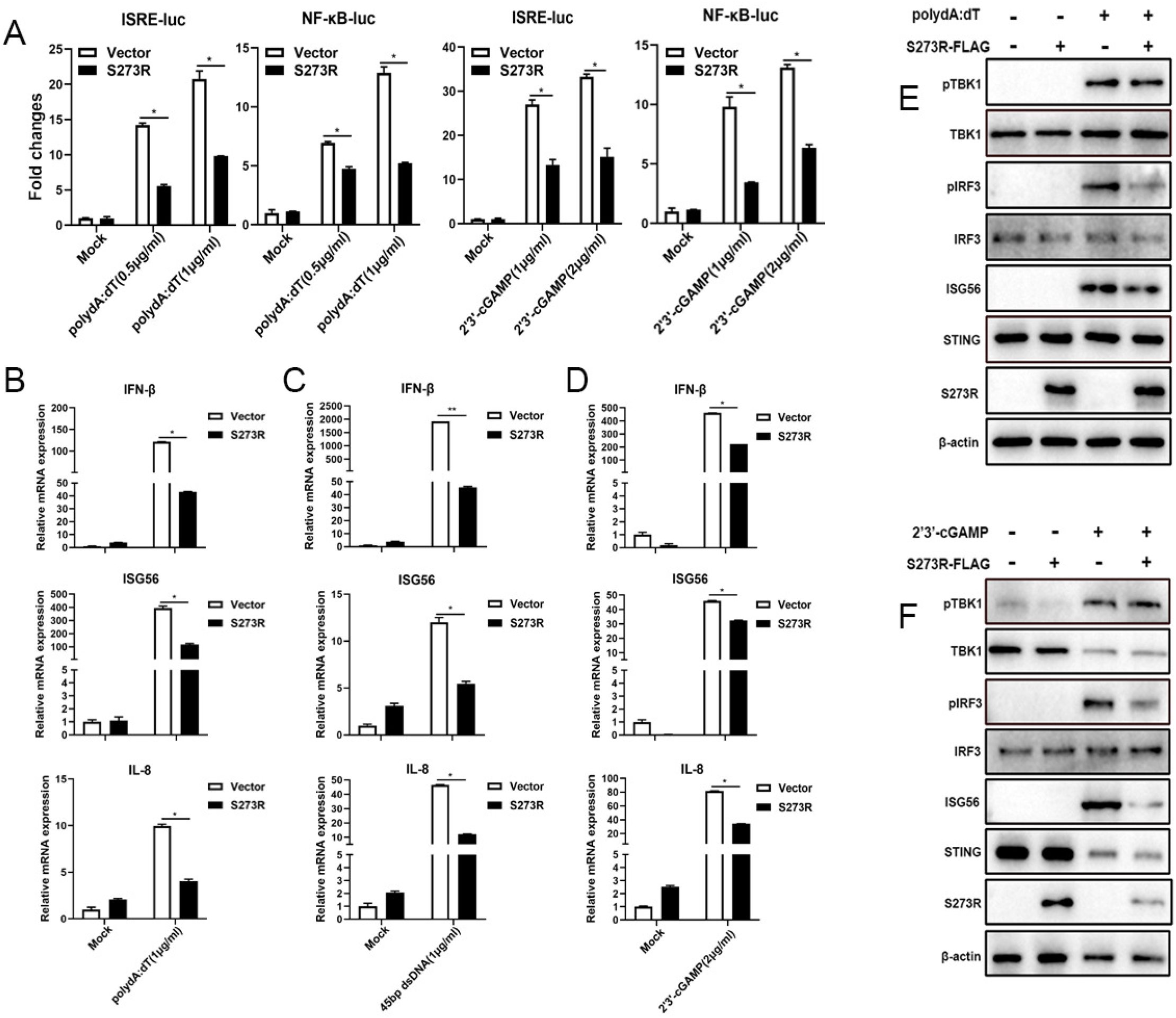
The effect of ASFV pS273R on endogenous porcine cGAS-STING signaling pathway. (A) PAMs in 96-well plates (2×10^4^ cells/well) were co-transfected with 10 ng ISRE-Luc or NF-κ?-Luc and 0.2 ng pRL-TK plasmids along with 20 ng 3×FLAG-pCMV-S273R or control vector 3 ×FLAG-pCMV, which were normalized to 50 ng/well by control vector. After 24h, the cells were not or transfected with polydA:dT (0.5μg/mL, 1μg/mL) or 2’3’-cGAMP (1μg/mL, 2μg/mL) for 8h. The luciferase activities were detected using Double-Luciferase Reporter Assay. (B-D) PAMs in 24-well plates (3×10^5^ cells/well) were co-transfected with 1μg 3×FLAG-pCMV-S273R or control 3×FLAG-pCMV for 24h and then transfected with polydA:dT (1μg/mL) (B) or 45bp dsDNA (1μg/mL) (C) or 2’3’-cGAMP (2μg/mL) (D) for 8h.The harvested cells were measured by RT-qPCR for downstream gene expressions as indicated. (E-F) PAMs in 24-well plate (3×10^5^ cells/well) were co-transfected with 1μg 3×FLAG-pCMV-S273R or control 3×FLAG-pCMV for 24h and then transfected with polydA:dT (1μg/mL) or 2’3’-cGAMP (5μg/mL) for 8h. The harvested cells were detected by Western blotting with the indicated antibodies.

### ASFV pS273R disturbs the porcine cGAS-STING signaling pathway mediated antiviral function

The innate immune cGAS-STING signaling pathway is capable of sensing virus attack and induces an antiviral response (19). To determine whether pS273R could disturb cGAS-STING mediated antiviral function, we selected two GFP viruses Herpes Simplex Virus-1 (HSV-1, a DNA virus) and Vesicular Stomatitis Virus (VSV, an RNA virus) to infect PAMs transfected with ASFV pS273R (Fig 3 and SupFig 2). The virus replications were examined by fluorescence microscopy, Western blotting, flow cytometry, RT-qPCR and TCID50 assay, respectively (Fig 3). We found that HSV-1 replicative GFP signals using MOI 0.01 and 0.1 were both enhanced in the presence of pS273R relative to controls under fluorescence microscopy (Fig 3A), by Western blotting (Fig 3B) and by flow cytometry analysis (Fig 3C). Using RT-qPCR, the HSV1 gB gene transcriptions were also upregulated with pS273R, in sharp contrast with the downregulated IFNβ and ISG56 genes (Fig 3D). Accordingly, the virus titers in the supernatants of PAMs infected with HSV1 of MOIs 0.01 and 0.1 were both increased relative to the controls in the TCID50 assay (Fig 3E).

**Figure 3.**
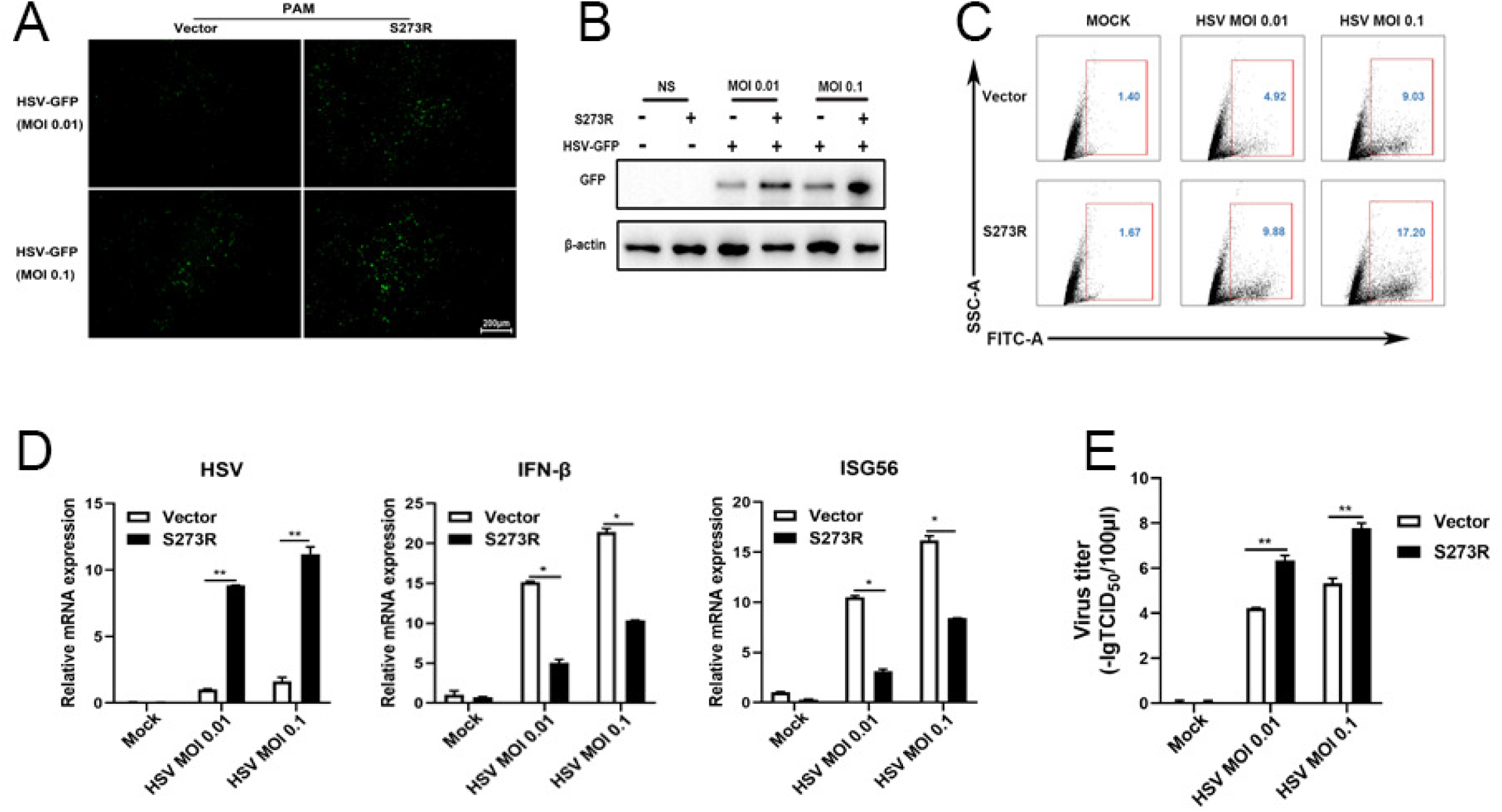
ASFV pS273R interfered the cGAS-STING signaling mediated anti-HSV-1 activity. (A-B) PAMs in 24-well plates (3×10^5^ cells/well) were co-transfected with 1μg 3×FLAG-pCMV-S273R or control 3×FLAG-pCMV for 24h and then infected with 0.01 MOI or 0.1 MOI HSV-1-GFP for 36 h, The GFP signals were observed by fluorescence microscopy (A). The infected cells were harvested to measure the GFP protein expressions by Western blotting (B). PAM cells in 6-well plates (8×10^5^ cells/well) were co-transfected with 2μg 3×FLAG-pCMV-S273R or control 3 × FLAG-pCMV for 24h and then infected with 0.01 MOI or 0.1 MOI HSV-1-GFP for 36 h, the GFP cells in infected PAMs were analyzed by flow cytometry. (D) The infected cells were harvested to measured HSV-1 gene and cellular gene transcriptions by RT-qPCR. (E) The viral titer in the supernatant from HSV-1 infected PAMs was measured by TCID50 assay.

The cGAS-STING pathway mediates a broad range of antiviral function including anti-DNA and anti-RNA activity, thus we also tested the role of pS273R during VSV replication (SupFig 2). Similarly, the VSV replicative GFP signals using MOIs 0.001 and 0.01 were both enhanced relative to the controls under fluorescence microscopy (SupFig 2A) and by flow cytometry (SupFig 2B). In RT-qPCR, the VSV glycoprotein gene transcriptions were upregulated with pS273R (SupFig 2C) in sharp contrast with the downregulated IFNβ and ISG56 gene transcriptions (SupFig 2E and 2F). Accordingly, the virus titers in the supernatants of infected PAMs with VSV of MOIs 0.001 and 0.01 were both increased with the S273R compared with the controls by the TCID50 assay (SupFig 2D) as well as by the plaque assay (SupFig 2G). Collectively, these results demonstrated that pS273R could enhanced virus replications by damaging cGAS-STING mediated antiviral functions.

### ASFV pS273R negatively regulates the cGAS-STING signaling pathway by targeting IKKε

To investigate the mechanism of how pS273R suppresses the cGAS-STING signaling pathway, pS273R was first co-transfected with individual signaling molecules of cGAS-STING-IFN pathway including TBK1 (Fig 4A), IKKε (Fig 4B) or IRF3-5D (Fig 4C) into 293T cells and the downstream ISRE, IFNβ and NF-κB promoter activity were examined. The results showed that only IKKε but not TBK1 and IRF3-5D activated promoter activity were inhibited by pS273R (Fig 4A-C). The cGAS-STING signaling also activates downstream NF-κB and pro-inflammatory cytokines even though the strength is much weaker and the associated molecular details are not clear. Thus, the signaling activity of NF-κB relevant molecules IKKβ and p65 were also checked in the presence of pS273R by NF-κB promoter assay, and it turned out that no any inhibitory effect of pS273R was observed (Fig 4D and 4E). Since only IKKε was affected by pS273R, the IKKε induced downstream gene transcriptions were examined by RT-qPCR, and the results showed that the IKKε induced downstream IFNβ, ISG56 and IL-8 gene levels were inhibited by pS273R in dose dependent manners (Fig 4F), which is consistent with the promoter assays (Fig 4B).

**Figure 4.**
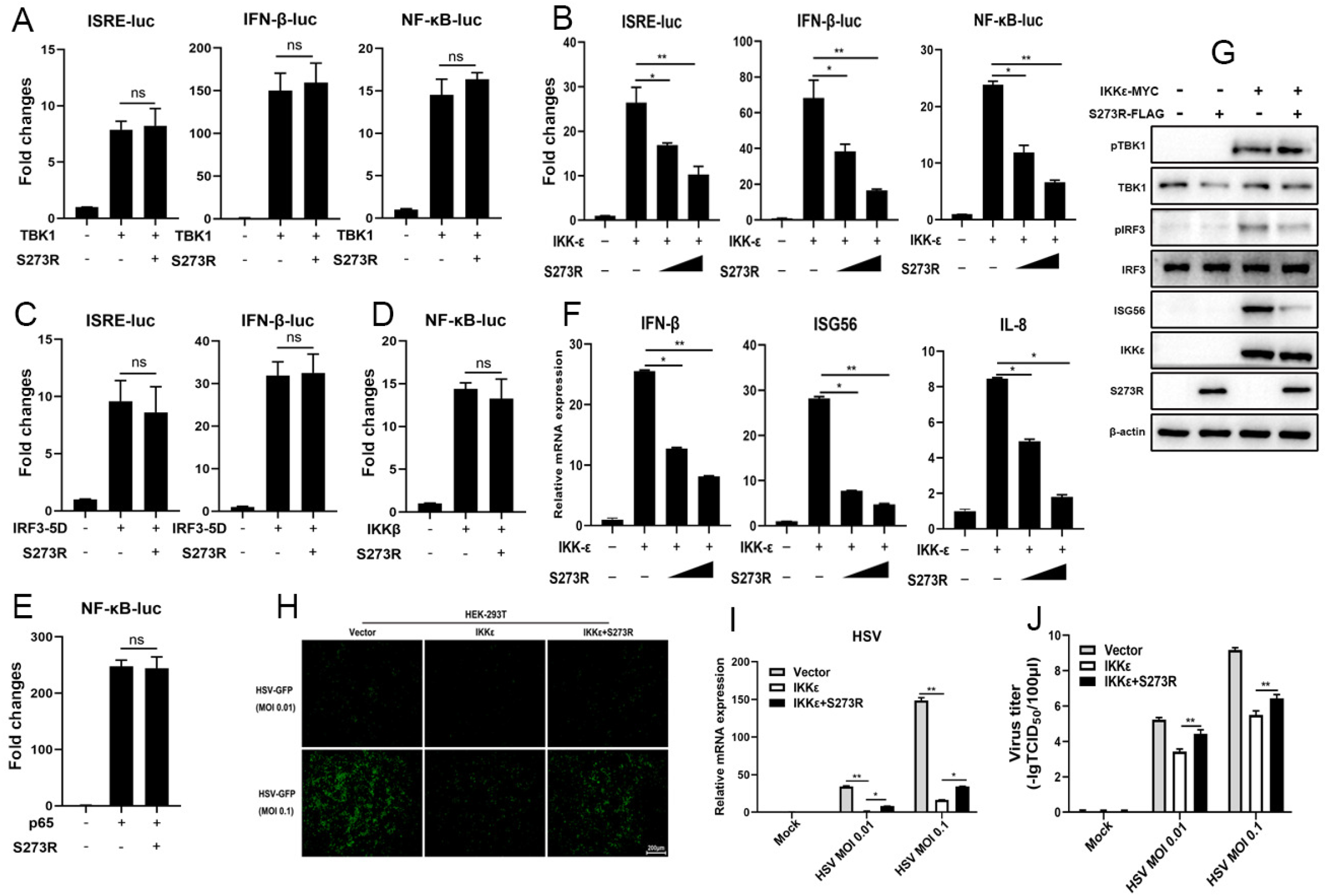
The identification of IKKε as the cellular target by ASFV pS273R. (A-E) HEK293T cells were co-transfected with 20 ng TBK1 expression plasmids and 10 ng 3×FLAG-pCMV-S273R (A), with 20 ng IKKε expression plasmid and increased amounts of 3×FLAG-pCMV-S273R (10 ng or 20 ng) (B), with 20 ng IRF3-5D expression plasmid and 10 ng pS273R (C), with 20 ng IKKβ plasmid and 10 ng pS273R (D), with p65 plasmid and 10 ng pS273R (E), plus 10 ng ISRE-Luc, IFNβ-Luc or NF-κB-Luc and 0.2 ng pRL-TK plasmids, which were normalized to 50 ng/well by vector 3×FLAG-pCMV After 24h, the luciferase activities were detected using Double-Luciferase Reporter Assay. (F-G) pS273R plasmid (400 ng or 800 ng) were co-transfected with 400 ng IKKε into HEK293T cells. After 24h, the cells were harvested and analyzed by RT-qPCR (F) and Western blotting (G). (H-J) The IKKε plasmid (500 ng) were co-transfected with 500 ng pS273R plasmid or empty vector into 293T cells for 24h, then the transfected 293T cells were infected with 0.01 MOI or 0.1 MOI HSV-1-GFP for 36 h, the GFP signals were observed by fluorescence microscopy (H). The infected cells were harvested to measure the HSV-1 gene expression by RT-qPCR (I). The viral titer in the supernatant from HSV-1 infected 293T cells was measured by TCID50 assay (J).

Next, the IKKε induced signaling was examined by Western blotting (Fig 4G). Ectopic IKKε was able to activate the phosphorylations of TBK1 and IRF3 and the downstream ISG56 production as previously reported (20). In the presence of pS273R, the IRF3 phosphorylation and ISG56 induction were inhibited, but once again TBK1 phosphorylation was not affected (Fig 4G). Finally, the IKKε induced antiviral activity was also investigated in the presence of pS273R (Fig 4H-J). IKKε exhibited obvious anti-HSV1 activity, however, in the presence of pS273R, the IKKε mediated anti-HSV1 activity was weakened as observed by fluorescence microscopy (Fig 4H), shown by RT-qPCR (Fig 4I) and TCID50 assay (Fig 4J). Similarly, pS273R promoted the VSV replications by damaging IKKε mediated antiviral function (SupFig 3). Collectively, these results illustrated that pS273R inhibited cGAS-STING signaling pathway by targeting IKKε.

### ASFV pS273R interacts with IKKε and disturbs the interaction between IKKε and STING

IKKε is a crucial IKK-related kinase that is significant for innate immune signaling, yet the interaction of STING and IKKε is still not clear. We first assessed the interaction between IKKε and STING by Co-IP, and the result showed that there was an obvious interaction between IKKε and STING (Fig 5A). Since pS273R targets IKKε and inhibits its activity, the interaction between pS273R and IKKε was also discernable in Co-IP assay (Fig 5C). However, there was no interaction between pS273R and STING in Co-IP assay (Fig 5B). Consistent results were shown for cellular co-localization between these proteins in PAMs by con-focal microscopy (Fig 5E). Obvious co-localization between IKKε and STING, co-localization between IKKε and pS273R, and no co-localization between pS273R and STING were observed (Fig 5E). The co-localization of STING and IKKε was so obvious that the punctation formation was appreciated (Fig 5E). However, in the Co-IP assay and in the presence of pS273R, the interaction between IKKε and STING was disturbed in a pS273R dose dependent manner (Fig 5D). These results indicated that ASFV pS273R blocks the interaction between STING and IKKε in a dose dependent manner to evade host innate immunity.

**Figure 5.**
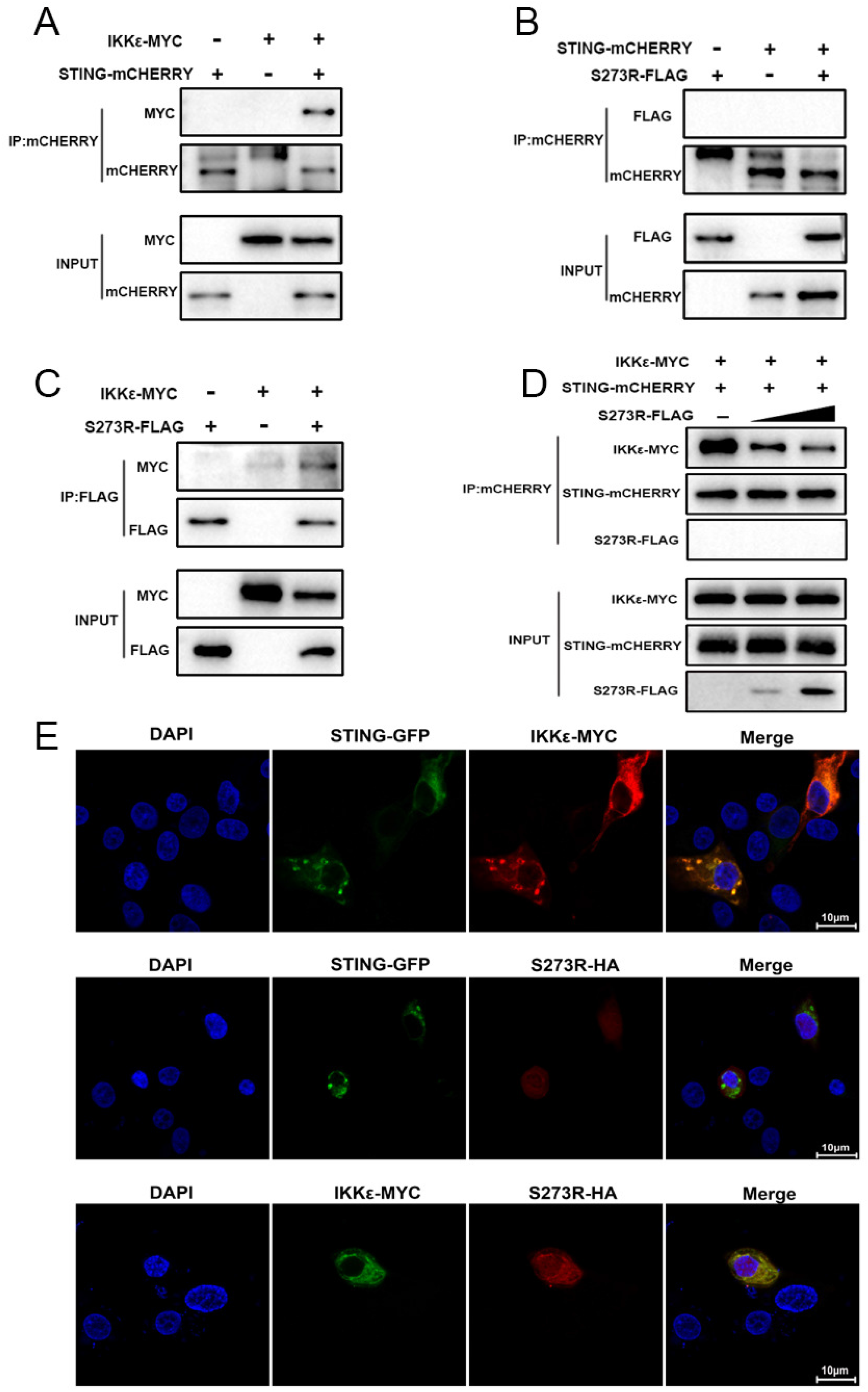
pS273R interacted with IKKε and disturbed the interaction between IKKε and STING. (A-C) HEK293T cells in 6-well plates (8×10^5^ cells/well) were co-transfected with 1μg each plasmid as indicated for 48h and then the cells were harvested and subjected for Co-IP using the indicated antibodies and subsequent Western blot analysis. (D) Increasing amounts of pS273R plasmid (1μg, 2μg) were co-transfected with 1μg STING-mCHERRY and 1μg IKKε-MYC into 293T cells for 48h and then cells were harvested and immunoprecipitated with anti-mCHERRY antibody and subjected to Western blot analysis. (E) PAM cells in 24-well plates (3×10^5^ cells/well) were co-transfected with STING-GFP (0.5μg) and IKKε-MYC (0.5μg), with STING-GFP (0.5μg) and pS273R-HA (0.5μg), with IKKε-MYC (0.5μg) and pS273R-HA (0.5μg) for 24h, and then cells were fixed and examined for cellular co-localization by con-focal microscopy.

### The protease activity of pS273R is responsible for inhibition of porcine cGAS-STING signaling pathway

A recent study about the structural information of pS273R revealed that a catalytic triad of C232-H168-N187 and the regions spanning amino acids 1-20 and 256-273 of pS273R play essential roles in pS273R enzyme activity (21). To explore the potential roles of these active sites of pS273R in the inhibitory function, we generated three point mutants (H168R, N187A and C232S) and two truncated mutants (ΔN1-20 and ΔN256-273) (Fig 6A). These enzyme inactive mutants were tested for inhibition of cGAS-STING signaling in various promoter assays. As shown in Fig 6B, the inhibitory effects of the five mutants disappeared at varying degrees. Furthermore, the inhibitory effects of the five mutants on IKKε activated promoter activity also disappeared (Fig 6C). To figure out whether these five mutants could disturb STING interaction with IKKε, the Co-IP of STING and IKKε was performed in the presence of each pS273R mutants, and the results showed that the pS273R mutants lost the ability to disturb the interaction of STING and IKKε (Fig 6D), with the mutant H168R exceptional, which reflected by the remained inhibition of cGAS-STING signaling (Fig 6B). The cellular co-localizations between STING and IKKε in the presence of pS273R mutants were examined and the results showed that with the pS273R, the co-localized punctuation patterns of STING and IKKε disappeared, whereas in the presence of mutants except H168R and ^Δ^N256-273, the significant co-localized punctuation patterns were maintained (SupFig 4).

**Figure 6.**
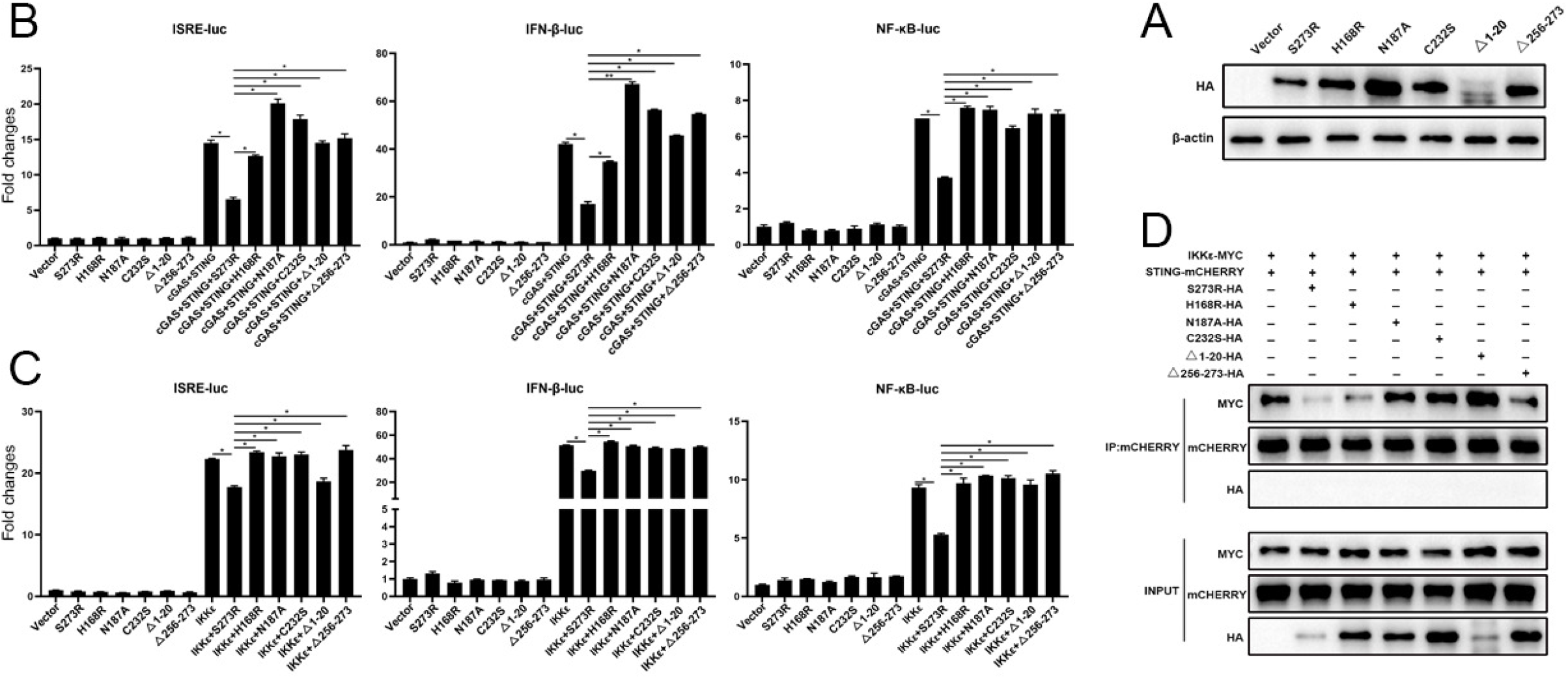
pS273R mutants without protease activity had decreased suppression on IKKε of cGAS-STING pathway. (A) HEK293T cells were co-transfected with 1μg of each pS273R mutants, respectively, and then the cells were harvested and analyzed by Western blotting. (B) HEK293T cells in 96-well plates (2×10^4^ cells/well) were co-transfected with 20 ng cGAS-HA and 10 ng STING-mCHERRY, plus 10 ng ISRE-luc or IFNβ-luc or NF-κB-luc and 0.2 ng pRL-TK plasmid, along with 10 ng pS273R or pS273R mutants, which were normalized to 50 ng/well by vector pCAGGS. At 24h post-transfection, luciferase activities were detected using Double-Luciferase Reporter Assay. (C) HEK293T cells were co-transfected with 20 ng IKKε, plus 10 ng ISRE-luc or IFNβ-luc or NF-κ?-luc and 0.2 ng pRL-TK plasmid, along with 10 ng pS273R or pS273R mutants, which were normalized to 50 ng/well by vector pCAGGS. After 24h, luciferase activities were measured. (D) Each pS273R mutants (1μg) were co-transfected with STING-mCHERRY (1μg) and IKKε-MYC(1μg) into 293T cells for 48h, and then the cells were harvested and subjected for Co-IP using anti-mCHERRY antibody and subsequent Western blot analysis.

It appeared that pS273R protease activity plays a critical role in the inhibition of cGAS-STING signaling as well as inhibition of IKKε activity. However, we did not observe the cleavage of IKKε and related signaling proteins, we wondered if pS273R, as a SUMO-1 protease, might affect the sumoylation modification of the target protein and its subsequent function. Therefore, we chose 2-D08, a unique inhibitor of SUMOylation (22, 23), to evaluate the hypothesis. The results showed that 2-D08 could markedly inhibit the IFNβ, ISG56 and IL-8 mRNA induction levels in a dose manner upon either polydA:dT stimulation (Fig 7A) or 2’3’-cGAMP stimulation (Fig 7B). The inhibitor 2-D08 also inhibited cGAS-STING signaling induced IFNβ and ISG56 mRNA levels in transfected 293T cells (SupFig 5). Analogously, 2-D08 was able to inhibit the interaction of IKKε and STING in Co-IP assay in a dose dependent manner (Fig 7C). Taken together, these data demonstrate that pS273R likely acts as an inhibitor of protein sumoylation to obstruct the interaction of IKKε and STING, and thus the cGAS-STING signaling and antiviral function.

**Figure 7.**
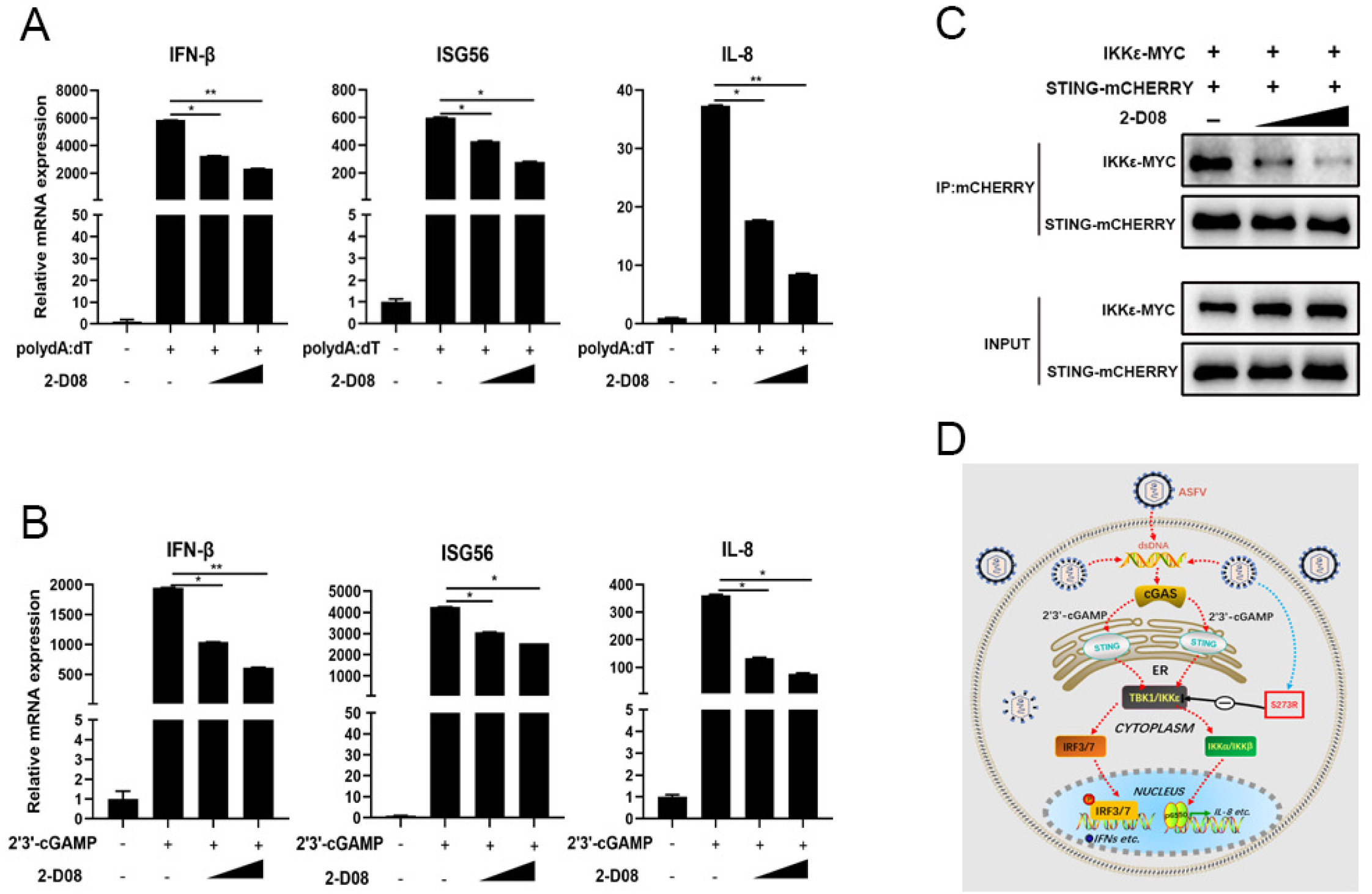
The inhibition of cGAS-STING signaling and interaction of STING and IKKε by the SUMOylation inhibitor. (A-B) PAMs in 24-well plates (3×10^5^ cells/well) were stimulated with polydA:dT (1μg/ml) (A) or 2’3’-cGAMP (2μg/ml) (B) for 8h and then treated with 2-D08 (50μM, 100μM) for l8h.The harvested cells were analyzed for downstream gene transcriptions by RT-qPCR. (C) HEK293T cells in 6-well plates (8×10^5^ cells/well) were co-transfected with STING-mCHERRY (1μg) and IKKε-MYC (1μg) for 24h and then treated with 2-D08 (50μM, 100μM) for 18h. The cells were harvested and subjected for Co-IP using anti-mCHERRY antibody and subsequent Western blot analysis. (D) The schematic diagram of ASFV pS273 antagonism on IKKε of cGAS-STING signaling pathway.

## Discussion

Many viruses are equipped to promote the replications and establish persistent infections in host. Therefore, viruses must evolved multiple mechanisms to antagonize the antiviral immune responses to achieve the immune evasion. Likewise, ASFV has developed multiple strategies to evade innate immune system, including modulation of type I IFN responses, inhibition of apoptosis, inhibition of autophagy and inhibition of inflammatory responses *etc*, with few of the corresponding viral proteins identified (24, 25). However, ASFV long double-stranded DNA genome codes for more than 150 proteins of which many are non-essential for viral replication in cells and their potential roles and the mechanism of action in evading the host’s defenses are still unknown. Here, we revealed for the first time that pS273R plays as a negative regulator in the cGAS-STING pathway via genomic ORF screening and confirming experiments. We also found that pS273R obviously promotes the replications of both DNA and RNA viruses, suggesting that pS273R facilitates their replications by damaging the cGAS-STING signaling which has been shown to have broad antiviral function (26).

Surprisingly, our results showed that pS273R inhibits the cGAS-STING signaling pathway by targeting and impairing the function of IKKε, suggesting that IKKε is likely to play an important role in the cGAS-STING pathway. Further, in the context of virus infections, pS273R has been observed to disturb IKKε mediated antiviral function. Therefore, our findings support that pS273R certainly acts on IKKε to impair the antiviral function and help virus evade host defense. IKKε has been previously reported to recruit and phosphorylate IRF3/7 in RIG-I-like receptors (RLR) and Toll-like receptors (TLR) signaling pathways (27, 28), whereas it has been long believed that TBK1 acts as the only downstream kinase to mediate the cGAS-STING pathway to activate IFN production. However, one recent study showed that IKKε synergizes redundantly with TBK1 to mediated STING activated NF-κB signaling and inflammatory cytokine production, on the other hand, IKKε also participates in STING activated IFN production even though in a much less degree (29). The phosphorylation of STING Ser365 (Ser366 in humans) is mediated by active TBK1, but the phosphorylation site of STING by IKKε in the absence of TBK1 is still not clear (29). Therefore, our findings indicate that IKKε is actually significant for the activation of the STING-dependent DNA sensing pathway, though the mechanism was unclear, which is warranted for future investigation.

Many viruses such as polio virus (30) and flavivirus (31) encode polyproteins or polyprotein precursors that are cleaved by viral proteases to form vital components of the virus particle, which are critical for virus maturation and infectivity. The ASFV pS273R protease, as a member of SUMO-1-specific protease family, could accurately cleave two polyproteins, pp220 and pp62, at Gly-Gly-Xaa sites to produce important proteins required for ASFV morphogenesis (10). Therefore, it is possible that pS273R protease inhibits the interaction of STING and IKKε by cleaving the IKKε. However, neither IKKε nor STING expression was changed in the presence of pS273R. The ASFV pS273R protease consists of arm domain and core domain. While the arm domain is unique, the core domain shares similar structure fold and active triad with those of other SUMO-1 cysteine proteases, which contain Ubl (ubiquitin-like protein)-specific proteases (Ulp) in yeast and sentrin-specific proteases (SENP) in mammals (32, 33). These proteases mainly catalyze the deconjugation of SUMOylated proteins; however, certain SUMO proteases are also critical for SUMO precursor maturation, thus indirectly affecting SUMO conjugation. As such, some viral proteases have the ability to deconjugate certain small protein modifications to interrupt host innate immune responses (34–35). It is also possible that pS273R protease could inhibit interaction of STING and IKKε by cleaving small protein modification such as sumoylation of IKKε. Indeed, the sumoylation inhibitor 2-D08 suppresses the interaction of STING and IKKε, cGAS-STING signaling and IKKε activity, certifying the function of pS273R protease in the cGAS-STING pathway. Collectively, our study presents clear evidence that pS273R protease acts as a sumoylation inhibitor and targets IKKε to interfere with the immune response.

In summary, upon infection of ASFV, porcine cytosolic cGAS senses the dsDNA of virus and catalyzes the synthesis of the second messenger cyclic GMP-AMP (2’3’-cGAMP), which binds and activates the adaptor STING. The oligomerized STING transfers from the endoplasmic reticulum (ER) to the trans-Golgi apparatus network (TGN); during the process, the TBK1 and IKKε as well are recruited, phosphorylated and activated. TBK1 and IKKε subsequently phosphorylate the IRF3/7 and activate IKKα/IKKβ, inducing expression of the interferons and inflammatory factors (Fig 7D). However, ASFV pS273R could antagonize the cGAS-STING signaling pathway by targeting the IKKε, thus achieving immune escape (Fig 7D).

## Materials and methods

### Cells and viruses

HEK-293T cells were cultured in Dulbecco modified Eagle medium (DMEM, Hyclone Laboratories, USA) supplemented with 100 IU/mL of penicillin plus 100 μg/mL streptomycin and 10% fetal bovine serum (FBS). Porcine alveolar macrophages (PAMs, 3D4/21) were cultured in RPMI 1640 medium (Hyclone Laboratories) which contains 100 IU/mL of penicillin plus 100 μg/mL streptomycin and 10% FBS. Cells were grown at 37°C in a 5% CO_2_ humidified incubator. The Vesicular Stomatitis Virus (VSV-GFP) and Herpes Simplex Virus-1 (HSV-1-GFP) were both provided by Dr. Tony Wang in SRI International USA.

### Reagents and antibodies

TRIpure Reagent for RNA extraction was purchased from Aidlab (Beijing, China). HiScript® 1st Strand cDNA Synthesis Kit, ChamQ Universal SYBR qPCR Master Mix, 2×Taq Master Mix (Dye plus), 180kDa prestained protein marker and TransDetect Double-Luciferase Reporter Assay Kit were all from Vazyme Biotech Co., Ltd (Nanjing, China). The Golden Star T6 Super PCR mix polymerase and KOD plus neo polymerase were from Tsingke (Nanjing, China) and Toyobo (Shanghai, China), respectively. The 2×MultiF Seamless Assembly Mix was acquired from Abclonal (Wuhan, China). Restriction endonucleases *Hind* III*, Kpn* I, *EcoR* I, *EcoR* V, *Dnp* I were purchased from New England Biolabs (Beijing, China). Agonists polydA:dT and 2’3’-cGAMP were bought from InvivoGen (Hongkong, China). Forty-five base pair double-stranded DNA (45bp dsDNA, tacagatctactagtgatctatgactgatctgtacatgatctaca) was synthesized by GENEWIZ (Shouzhou, China). Lipofectamine^TM^ 2000 were acquired from ThermoFisher Scientific (Shanghai, China). Chemical inhibitor 2-D08 was purchased from Selleck (Shanghai, China). Protein A/G Plus-Agarose was from Santa Cruz Biotechnology (Dallas, Texas, USA). Mouse anti-FLAG mAb, mouse anti-Actin mAb, mouse anti-GFP mAb were all acquired from Transgen Biotech (Beijing, China). The HRP anti-mouse IgG, HRP anti-rabbit IgG were purchased from Sangon Biotech (Shanghai, China). The ColorMixed Protein Marker was bought from Solarbio Biotech (Beijing, China). The rabbit anti-TBK1 (D1B4), anti-p-TBK1 (S172), anti-p-IRF3 (4D4G), anti-IRF3 (D164C), anti-HA (C29F4) were bought from Cell Signaling Technology (Danvers, MA, US). Rabbit anti-STING pAb was from ProteinTech (Wuhan, China). Rabbit anti-ISG56-pAb was homemade in our lab. DAPI staining solution was bought from Beyotime Biotech (Shanghai, China). Rabbit anti-mCHERRY-pAb, mouse anti-MYC mAb, Goat anti-mouse IgG H&L (Alexa Fluor®594) and goat anti-rabbit IgG H&L (Alexa Fluor®647) were purchased from Abcam (Shanghai, China).

### Plasmid construction and gene mutations

The genomic ORFs of ASFV China 2018/1 (GenBank submission No: MH766894) were codon optimized, synthesized and cloned into p3×FLAG-CMV-7.1 vector using *Not* I and *Sal* I sites. The 145 sequence confirmed ORF plasmids were used for screening of modulators of porcine cGAS-STING signaling pathway by ISRE promoter assay (Supplementary Fig 1). The S273R gene was amplified by PCR using Golden Star T6 Super PCR Mix polymerase from plasmid p3 × FLAG-CMV-S273R and then was cloned into pCAGGS-HA vector using *EcoR* I and *EcoR* V sites. The critical enzyme active sites of pS273R were identified and point mutated using the mutation PCR primers designed by QuickChange Primer Design method (https://www.agilent.com) (Supplementary Table 1). The mutation PCR was performed with KOD plus neo polymerase and pCAGGS-S273R-HA as the template. The PCR products were transformed into competent DMT *E. coli* after *Dpn* I digestion, and the resultant mutants were sequence confirmed. Truncated mutants of the pS273R, including pS273R (Δ1-20) and pS273R (Δ256-273) were obtained by PCR amplification and cloning into pCAGGS-HA using 2 × MultiF Seamless Assembly Mix polymerase. Porcine cGAS, STING, IKKε, IKKβ and NF-kB p65 plasmids were constructed and characterized as we reported (36); human TBK1, IKKε, IRF3-5D plasmids were preserved in our lab.

### Promoter driven luciferase reporter gene assay

293T cells grown in 96-well plates (3-4×10^4^ cells / well) were co-transfected by Lipofectamine 2000 (Thermo Fisher Scientific, Waltham, MA USA) with reporter plasmids, ISRE-Luc, IFNβ-Luc or ELAM (NF-κB)-firefly luciferase (Fluc) (10 ng/well) plus *Renilla* luciferase (Rluc) reporter (0.2 ng/well), with or without the indicated porcine cGAS and STING plasmids as well as porcine S273R or vector control (10-20 ng/well). The total DNA amount for transfection was normalized with control vectors to 50 ng for each well. PAMs grown in 96-well plates were similarly transfected using the Lipofectamine 2000. At 24h post transfection, cells were harvested and dual luciferase assays were sequentially performed using the TransDetect Double-Luciferase Reporter Assay Kit. The fold changes were calculated relative to control samples after normalization of Fluc by Rluc.

### Quantitative real-time PCR

Total RNA was extracted from 293T or PAMs in 24-well plates (2-4×10^5^ cells / well) with TRIpure Reagent, and cDNA synthesis was performed using HiScript® 1st Strand cDNA Synthesis Kit. The quantitative PCR was then performed with ChamQ Universal SYBR qPCR Master Mix using StepOne Plus equipment (Applied Biosystems) to measure the target gene expressions. The qPCR program is denaturing at 95 °C for 30 s followed by 40 cycles of 95 °C for 10 s and 60 °C for 30 s. The qPCR primers for hIFNβ, hISG56, hIL8, hRPL32, pIFNβ, pISG56, pIL1β, pIL8, pβ-actin, HSV-1 gB, VSV Glycoprotein and GFP are shown in Supplementary Table 2.

### Flow Cytometry and virus TCID50 assay

PAMs in 6-well plate (8×10^5^ cells/well) were transfected by Lipofectamine 2000 with porcine S273R expressing plasmids or vector control (1μg /well). About 24h post-transfection, the transfected cells were infected or not with VSV-GFP (0.001 MOI) or HSV-GFP (0.01 MOI). After infection, the cells were harvested by trypsin digestion and washed 3×with PBS. The levels of GFP virus replication were analyzed by flow cytometry of the cell suspensions.

The supernatants of virus infected PAMs were collected and tenfold serially diluted in DMEM medium with each dilution four to eight replications. The diluted supernatants were then used to infect Vero cells in 96-well plates for 2h. The cells were then cultured in DMEM containing 2% FBS and grown at 37°C for 1 day (for VSV) or 2 day for (HSV-1). The cytopathic effects (CPEs) were counted and TCID50 was calculated with Reed-Muench method.

### Co-immunoprecipitation and Western blot analysis

For Co-immunoprecipitation, 293T cells in 6-well plate (8×10^5^ cells/well) were transfected for 48h, and then cells were harvested and lysed in 500 μL RIPA buffer (50 mM Tris pH 7.2, 150 mM NaCl, 1% sodium deoxycholate, 1% Triton X-100) with protease inhibitors on ice for 30 min followed by centrifugation. The 50 μL cell lysate was saved as input controls and the remained were incubated with indicated antibodies overnight at 4 °C, and then 30 μL 50% protein A/G bead solution was added for another 2h. Later, the beads were thoroughly washed five times with RIPA and boiled in 40 μL 2×SDS sample buffer to obtain the elution which subjected to Western blotting. The elution samples together with input controls were boiled at 100 °C for 5-10 min, separated by 6-10% SDS-polyacrylamide gels, and transferred to PVDF membranes. Membranes were then blocked using 5% nonfat dry milk Tris-buffered saline, with 0.1% Tween-20 (TBST) at room temperature (RT) for 2h. Next the membranes were sequentially incubated with primary antibodies overnight at 4 °C and HRP-conjugated goat anti-mouse or anti-rabbit IgG for 1h at RT. Protein signals was visualized and captured by Western blot imaging system (Tanon, Shanghai, China).

### Confocal fluorescence microscopy

PAMs grown on coverslip in 24-well plates (2×10^5^ cells/well) were co-transfected by Lipofectamine 2000 with the combinations of STING-GFP, IKKε-MYC and S273R. About 24h post transfection, the cells were fixed with 4% paraformaldehyde at RT for 30 min and permeabilized with 0.5% Triton X-100 for 20 min. The cells were blocked using 2% bovine serum albumin in PBS at RT for 30 min, and then were incubated with primary anti-FLAG mouse mAb or anti-MYC mouse mAb (1:200) and primary anti-HA rabbit pAb (1:200) overnight at 4 °C. Next, cells were washed 3× in PBST (1-2 min per wash) and incubated with secondary antibodies Goat Anti-Mouse IgG H&L Alexa Fluor 594 (1:500) and Goat anti-Rabbit IgG H&L Alexa Fluor 647 (1:500). The cell nuclei were counter stained with 0.5μg/mL 4’,6-diamidino-2-phenylindole (DAPI, Beyotime, China) at 37°C for 15 min. Finally, the coverslip with stained cells was loaded on slide, sealed by nail polish, and visualized by laser-scanning confocal microscope (LSCM, Leica SP8, Solms, Germany) at the excitation wavelengths 405 nm, 488 nm, 561 nm and 633 nm, respectively.

### Statistical analysis

The data were the representative of two or three similar experiments and shown as the mean ± SD. The statistical significance was determined by analysis with the software GraphPad Prism 8.0, where *p* < 0.05 was considered statistically significant as determined by the student *t*-test. In the figures, “*”, “**” and “ns” denote *p* < 0.05, *p* < 0.01 and statistically not significant, respectively.

---

**Supplementary Figure 1.**
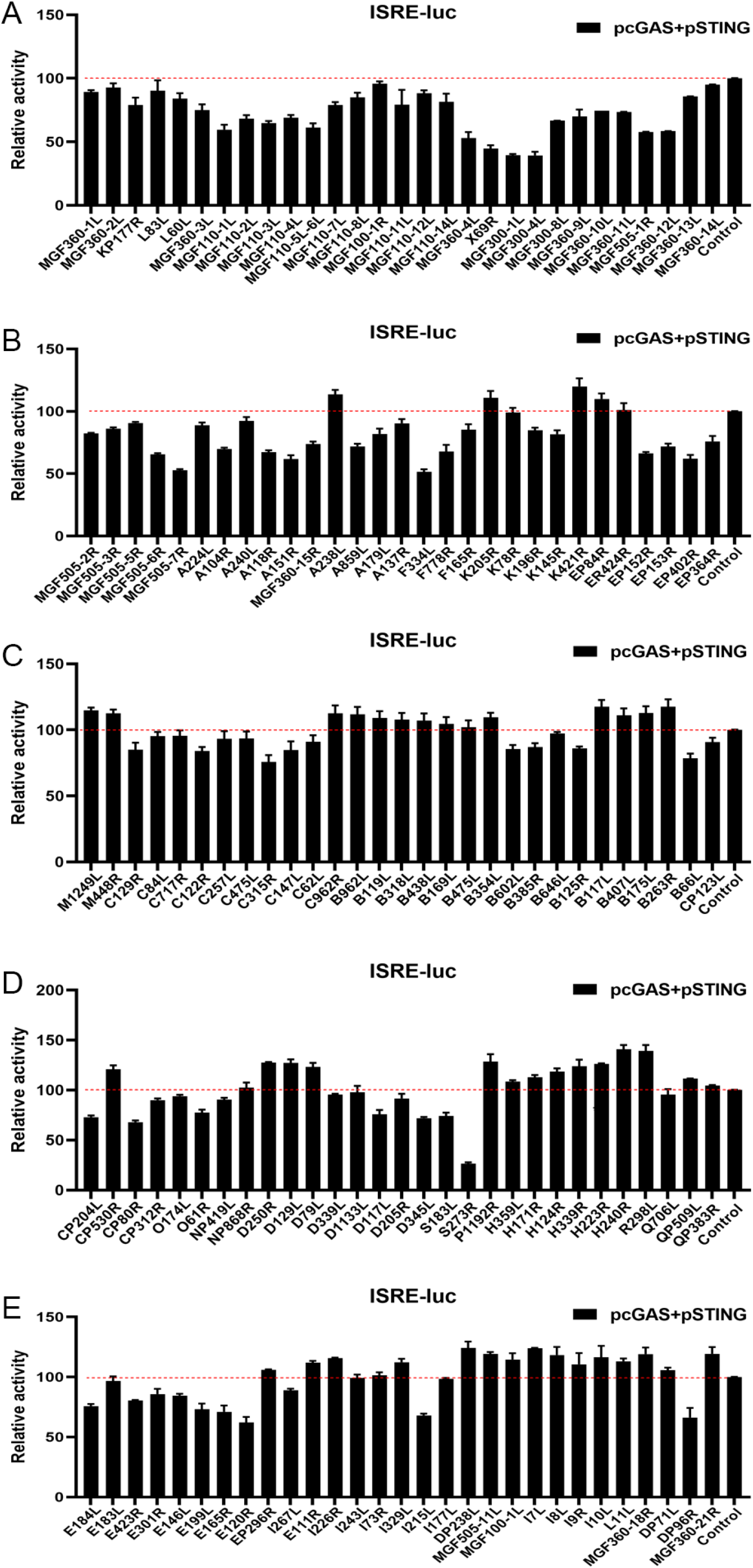
The screening of ASFV genomic ORFs for the modulation of porcine cGAS-STING signaling by ISRE promoter assay in transfected cells. (A) HEK293T cells in 96-well plates (2×10^4^ cells/well) were co-transfected with 20 ng cGAS-HA and 10 ng STING-GFP, plus 10 ng ISRE promoter plasmid and 0.2 ng pRL-TK plasmid, along with 10 ng ASFV ORF genes or empty vector 3×FLAG-pCMV, which were normalized to 50 ng/well by vector 3× FLAG-pCMV. At 24h post-transfection, luciferase activities were detected using Double-Luciferase Reporter Assay.

**Supplementary Figure 2.**
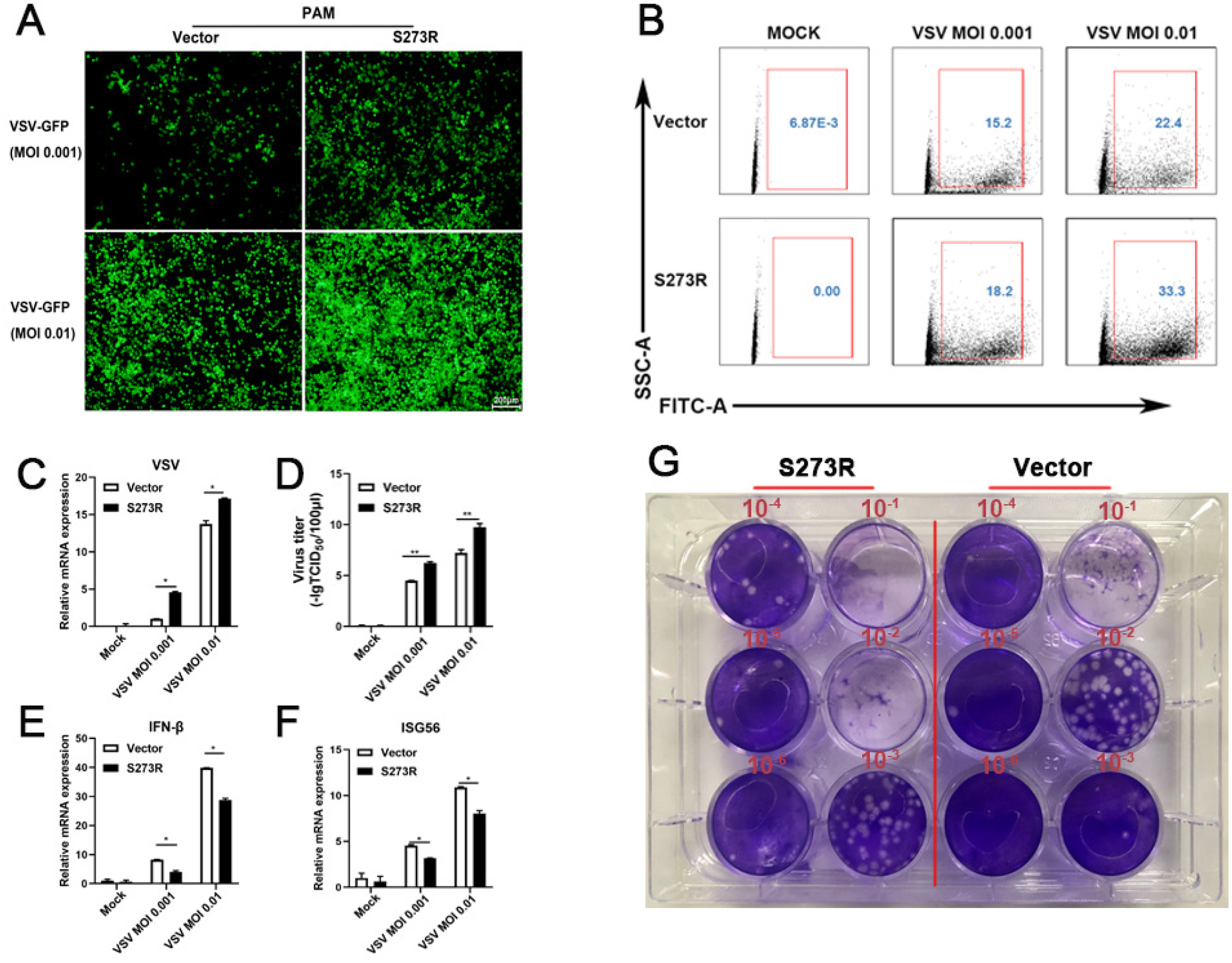
pS273R inhibited cGAS-STING signaling mediated anti-VSV function. (A) PAMs in 24-well plates (3×10^5^ cells/well) were co-transfected with 1μg 3×FLAG-pCMV-S273R or empty control 3×FLAG-pCMV for 24h and then infected with 0.001 MOI or 0.01 MOI VSV for 16 h. The GFP signals were observed by fluorescence microscopy. (B) PAM cells in 6-well plates (8×10^5^ cells/well) were co-transfected with 2μg 3×FLAG-pCMV-S273R or control 3× FLAG-pCMV for 24h and then infected with 0.001 MOI or 0.01 MOI HSV-1 for 16h, the GFP cells in infected PAMs were analyzed by flow cytometry. The infected cells were harvested to measured VSV-1 gene (C) and cellular gene transcriptions (E-F) by RT-qPCR. The viral titer in the supernatant from VSV-1 infected PAMs was measured by TCID50 assay (D) and plaque assay (G).

**Supplementary Figure 3.**
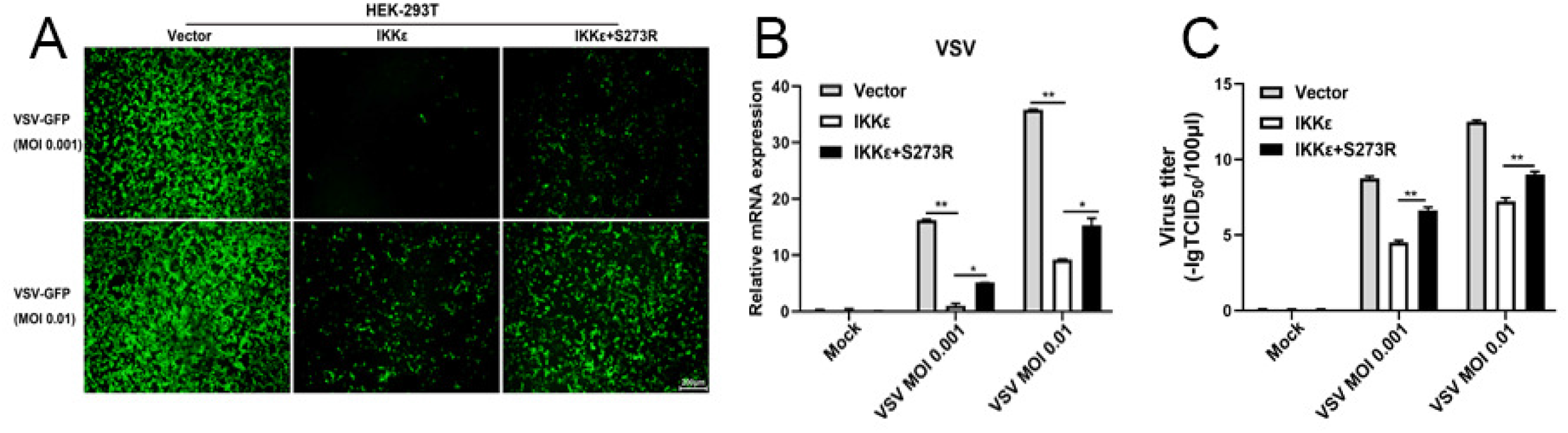
pS273R interfered the anti-VSV activity by IKKε. The IKKε (500 ng) were co-transfected with 500 ng pS273R plasmid or control vector into 293T cells for 24h, then the cells were infected with 0.001 MOI or 0.01 MOI VSV for 16h, the GFP signals were observed by fluorescence microscopy (A). The infected cells were harvested to measure the VSV gene expression by RT-qPCR (B). The viral titer in the supernatant from VSV infected 293T cells was measured by TCID50 assay (C).

**Supplementary Figure 4.**
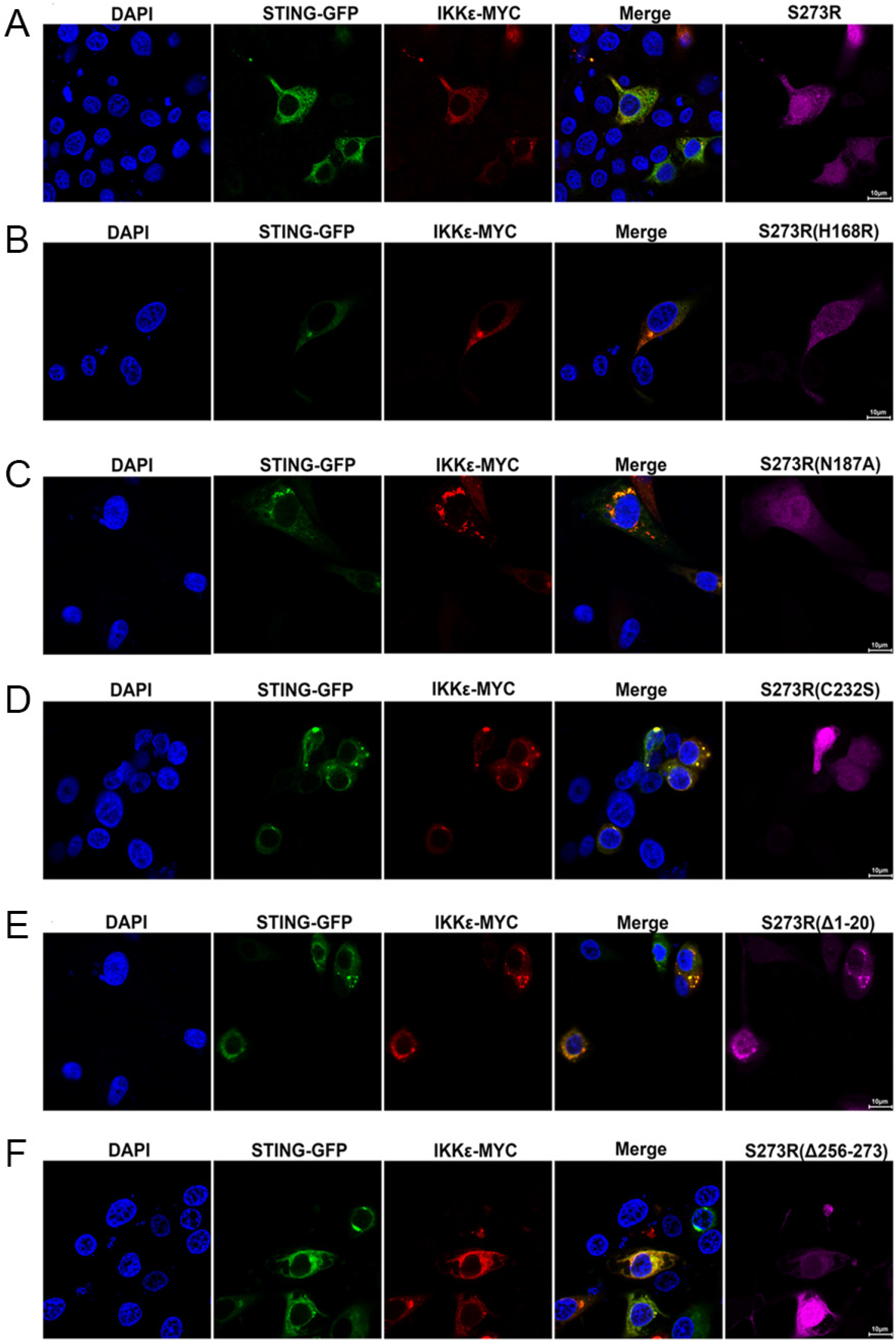
The cellular co-localizations between STING and IKKε in the presence of S273R and its various mutants. 0.4μg of each pS273R mutants or pS273R were co-transfected with STING-GFP (0.4μg) and IKKε-MYC (0.4μg) into PAMs for 24h, and then the cells were fixed, stained and examined for cellular co-localization by con-focal microscopy.

**Supplementary Figure 5.**
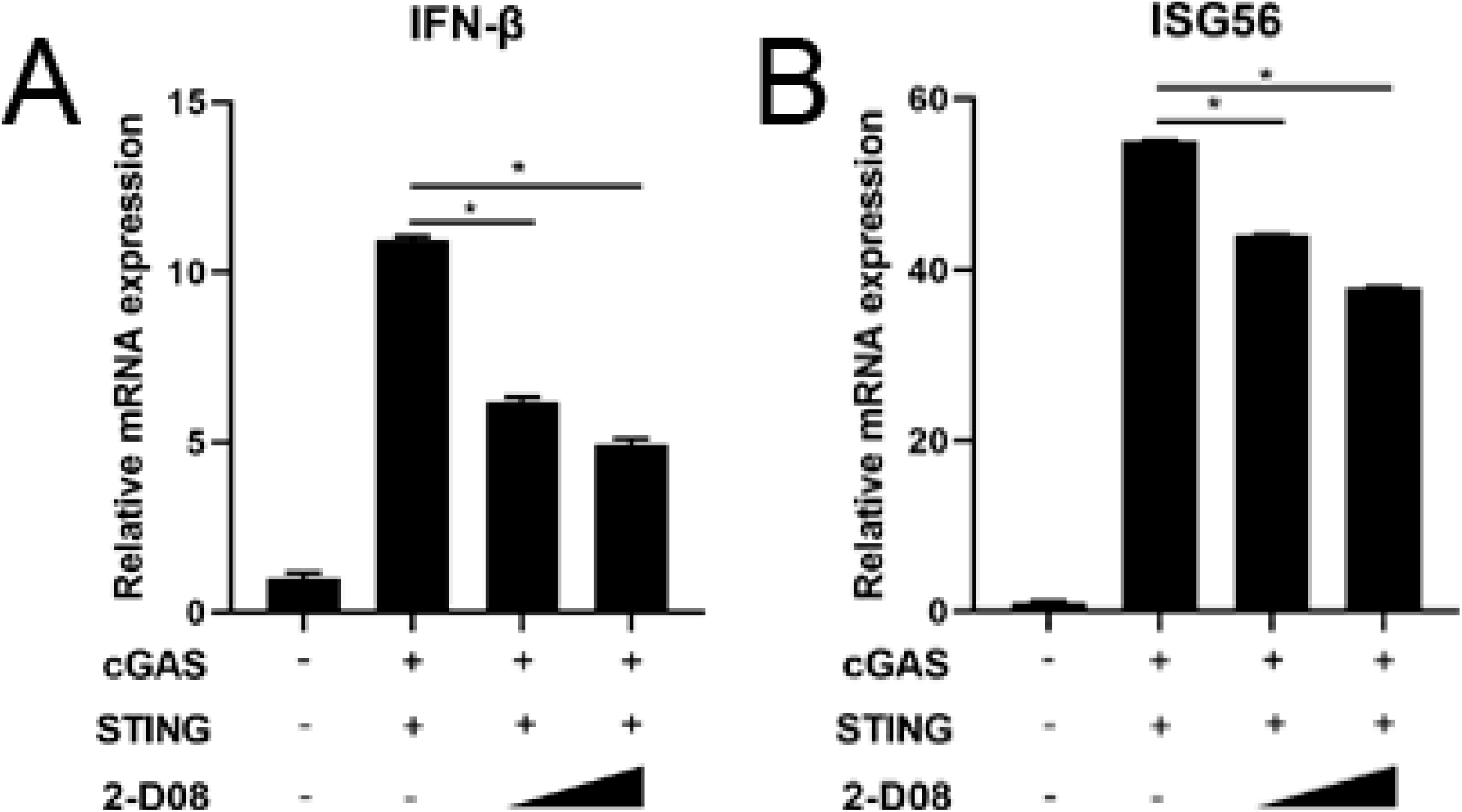
The inhibition of exogenous cGAS-STING signaling mediated downstream gene inductions by 2-D08. (A-B) HEK293T cells in 24-well plates (3×10^5^ cells/well) were co-transfected with cGAS-HA (0.5μg) and STING-GFP (0.5μg) for 24h, and then treated with 2-D08 (50μM, 100μM) for another 18h. The cells were harvested and subjected for analysis by RT-qPCR.

## Author contribution statement

J.Z conceived and designed the experiments; J.L, J.N, S.J, N.X, Y.G, J.Z, Q.C, performed the experiments; W.Z, N.C, Q.Z, H.C, X.G, H.Z, F.M, J.Z analyzed the data; J.L and J.Z wrote the paper. All authors contributed to the article and approved the submitted version.

## Acknowledgments

The work was partly supported by the National Key Research and Development Program of China (2017YFD0502301), Jiangsu provincial key R & D plan (BE2020398), the National Natural Science Foundation of China (31872450), and A Project Funded by the Priority Academic Program Development of Jiangsu Higher Education Institutions (PAPD).

